# Evidence for the Involvement of Vernalization-related Genes in the Regulation of Cold-induced Ripening in ‘D’Anjou’ and ‘Bartlett’ Pear Fruit

**DOI:** 10.1101/851733

**Authors:** Seanna Hewitt, Christopher A. Hendrickson, Amit Dhingra

## Abstract

European pear (*Pyrus communis L.*) cultivars require a genetically pre-determined duration of cold-temperature exposure to induce autocatalytic system 2 ethylene biosynthesis and subsequent fruit ripening. The physiological responses of pear to cold-temperature-induced ripening have been well characterized, but the molecular mechanisms underlying this phenomenon continue to be elucidated. This study employed established cold temperature conditioning treatments for ripening of two pear cultivars, ‘D’Anjou’ and ‘Bartlett’. Using a time-course transcriptomics approach, global gene expression responses of each cultivar were assessed at four different developmental stages during the cold conditioning process. Differential expression, functional annotation, and gene ontology enrichment analyses were performed. Interestingly, evidence for the involvement of cold-induced, vernalization-related genes and repressors of endodormancy release was found. These genes have not previously been described to play a role in fruit during the ripening transition. The resulting data provide insight into cultivar-specific mechanisms of cold-induced transcriptional regulation of ripening in European pear, as well as a unique comparative analysis of the two cultivars with very different cold conditioning requirements.

## Introduction

Pear (*Pyrus spp.*) is an economically important and nutritionally valuable tree fruit genus worldwide. European pear (*Pyrus communis L.*) cultivars are among the most widespread, commercially grown Pyrus members, and are cultivated in Europe, North America, South America, Africa, and Australia [1]. Along with apples, bananas, peaches, tomatoes, mangoes, and avocados, European pear is classified as climacteric in its ripening profile. Ripening in most climacteric fruits involves a seamless transition between system 1 (S1) and system 2 (S2) ethylene production. This is the point at which regulation of ethylene synthesis changes from autoinhibitory to auto-stimulatory [2, 3]. Increased ethylene biosynthesis during climacteric ripening is accompanied by a concomitant spike in respiration [2, 4, 5]. A prominent aspect that distinguishes European pear and Chinese white pear (*Pyrus bretschneideri* Rehder.) from most other climacteric fruits is that these species have a pre-ripening period during which they require a specific amount of cold exposure in order to transition from S1 to S2 ethylene production [6] [7]. The process of cold exposure is called ‘conditioning’ and accumulation of chilling hours necessary for this transition, and therefore ripening initiation, is referred to as the ‘chilling requirement’ [8].

The chilling requirement for ripening varies by cultivar. ‘Bartlett’ pears require 15 days of chilling, ‘Comice’ require 30, and ‘D’Anjou’ require 60 days of chilling at 0°C. ‘Passe Crassane’ pears lie at the extreme end of the conditioning spectrum, with a requirement of 90 days of chilling at 0°C to ripen. However, the duration of chilling may be manipulated by increasing the temperature at which conditioning is conducted, with an appropriate temperature range of 0-15°C [8]. In addition to genetic predetermination for chilling duration, chilling time is affected by maturity of the fruit at harvest, with pears harvested at a greater maturity index requiring a less extensive period of cold conditioning, and vice versa [9].

Physiological studies have characterized the conditioning requirements for a range of European pear cultivars under defined conditioning temperatures, exogenous ethylene application regimes, and other pre-harvest treatments [8–11]. While exogenous ethylene treatment reduces cold conditioning needs, in most European pear cultivars it does not entirely compensate for the need of cold conditioning. This indicates that cold-dependent mechanisms are partly responsible for regulating the development of ripening competency, which in turn impacts the quality and marketability of pear fruit. Interestingly, in contrast to *P. communis* and *P. bretschneideri*, many Japanese pear (*Pyrus pyrifolia L.*) varieties have no conditioning requirements and are also regarded as non-climacteric fruits because they do not display the characteristic S1 to S2 transition during ripening [12]. Additionally, ‘Bosc’, unique from other European pear cultivars, acquires competency for ripening with exogenous ethylene only, needing no chilling to ripen [8].

Requirement of cold exposure in pear to induce ripening is reminiscent of other natural cold temperature-dependent developmental phenomena, such as the vernalization and stratification that are needed for flowering and seed germination, respectively. The genes that regulate vernalization and stratification have been well-studied in model organisms, including Arabidopsis, wheat, and barley [13–15]; however, similar gene homologues have not yet been reported in cold-induced fruit ripening. With respect to flowering, the process of developmental initiation following an environmentally governed dormant state is known as endodormancy release [16]. Thus far, endodormancy release has not been used to characterize ripening after chilling, although the two processes share many similarities with regards to the timing and environmental nature of cold required, suggesting that similar genetic and regulatory mechanisms may govern these processes. Various forms of the chilling requirement for ripening have been described in avocado and mango, although to a lesser degree than in pear, as the former are more prone to chilling injury [17, 18].

Few recent studies have utilized a transcriptomics approach to characterize the molecular underpinnings of cold-induced S1 to S2 transition in Pear. There is a complex interaction of genes involved in regulating phytohormones, secondary messengers, signaling pathways, respiration and chromatin modification that underlie cold-induced progression of ripening [19–24]. Genes associated with phytohormones such as abscisic acid (ABA), auxin, and jasmonic acid along with transcription factors were implicated in low-temperature-mediated enhancement of ripening in ‘Bartlett’ [22]. In ‘Passe Crassane’ the impact of low temperature (LT) induced ethylene and exogenous ethylene treatments were evaluated using a transcriptomics approach [25]. It was observed that the expression of a subset of the low temperature-induced differentially expressed genes was disrupted by 1-MCP treatment indicating that they were regulated by LT-induced ethylene. It was also reported that several transcription factors were unaffected by 1-MCP treatment, implying that they were under the control of LT alone [25]. Recent work quantifying expression of key genes representing ripening-related metabolic pathways in ‘D’Anjou’ and ‘Bartlett’ pear cultivars during the process of cold-conditioning, revealed an increased alternative oxidase (AOX) expression prior to the onset of the ripening climacteric. This novel finding suggests that AOX may play an important role in the achievement of ripening competency [26] and may be necessary for the onset of the ripening climacteric in pear and other chilling dependent fruit. The Alternative Oxidase (AOX) respiratory pathway is known to play a role in cold stress mediation and response in many plant systems including cold temperature induced activation of respiration in potatoes, cell expansion and elongation in cotton, and mitigation of chilling injury in tomato and chickpea [27–29]. In many plant systems, AOX activity serves to maintain carbon metabolism homeostasis, cellular redox state, and ROS homeostasis during development [30].

Based on our previous work [26], a time course RNAseq analysis was performed in this study using ‘Bartlett’ and ‘D’Anjou’ fruit to evaluate the hypothesis that AOX and other key cold-induced genes facilitate ripening, and that the fruit from the two cultivars recruit different set of genes during cold conditioning. Key genes and networks involved in cold-induced, ripening-associated biochemical pathways were identified.

## Results and Discussion

### Fruit Firmness

Tissues from the fruit used in Hendrickson, Hewitt (31) were utilized for RNAseq analysis conducted in this study. Cold conditioning of the fruit at 10°C resulted in a reduction of fruit firmness in both ‘Bartlett’ and ‘D’Anjou’ cultivars as demonstrated (see Figure 2 in [31]). For both cultivars, fruit softening accelerated once the fruit was transferred to 20°C. The rate of softening was more rapid for ‘Bartlett’ than ‘D’Anjou’.

**Table 1.**
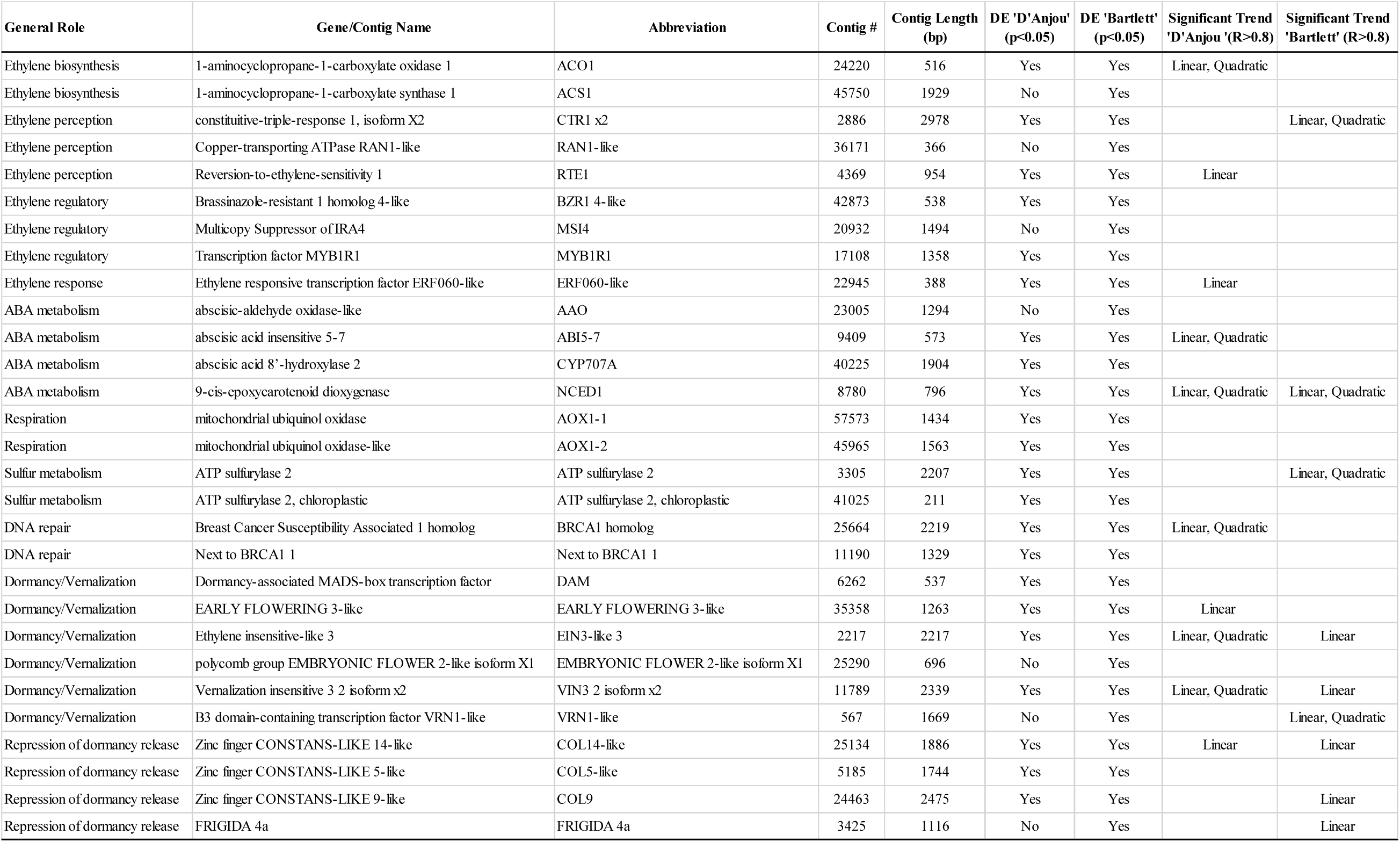
Summary of differentially expressed contigs. Information includes general role or associated pathway, full and abbreviated names, contig number (corresponding to sequences, annotations, and expression values in Supplementary Files 1 and 2), length, and indication of significant differential expression and/or significant expression trends.

### RNAseq Assembly Analysis

RNAseq assembly generated 140077 contigs (Supplementary File 1). In the OmicsBox suite, the maSigPro R package was used to conduct time course differential expression analyses for both cultivars. 17,711 differentially expressed contigs (p<0.05) were identified for ‘D’Anjou’, with 7,174 of these contigs exhibiting significant linear or quadratic trends over time (R>0.8). In ‘Bartlett’ 31,481 contigs were identified as being differentially expressed, with 7,174 contigs exhibiting significant quadratic or linear trends over time (R>0.8) (Supplementary File 2). Similarities and differences in expression trends of contigs of interest between ‘D’Anjou’ and ‘Bartlett’, as well as expression patterns of differentially expressed contigs (DECs) associated with ethylene and phytohormone metabolism, abscisic acid metabolism, TCA cycle, respiration, were assessed. Additionally, in order to better understand the mechanisms underlying the chilling requirement for ripening in *Pyrus*, the expression of genes associated with: vernalization, flowering, dormancy, and other processes directly induced by cold/chilling were observed. Pre-climacteric expression of *AOX1* peaked during conditioning prior to onset of ripening (Figure 1), supporting the hypothesis. Additional key genes, including those that mediate vernalization and endodormancy release were also observed to be differentially expressed. Detailed analysis of RNAseq results and enriched gene ontologies related to phytohormone metabolism and cold-response pathways are discussed in detail in the following sections.

**Figure 1.**
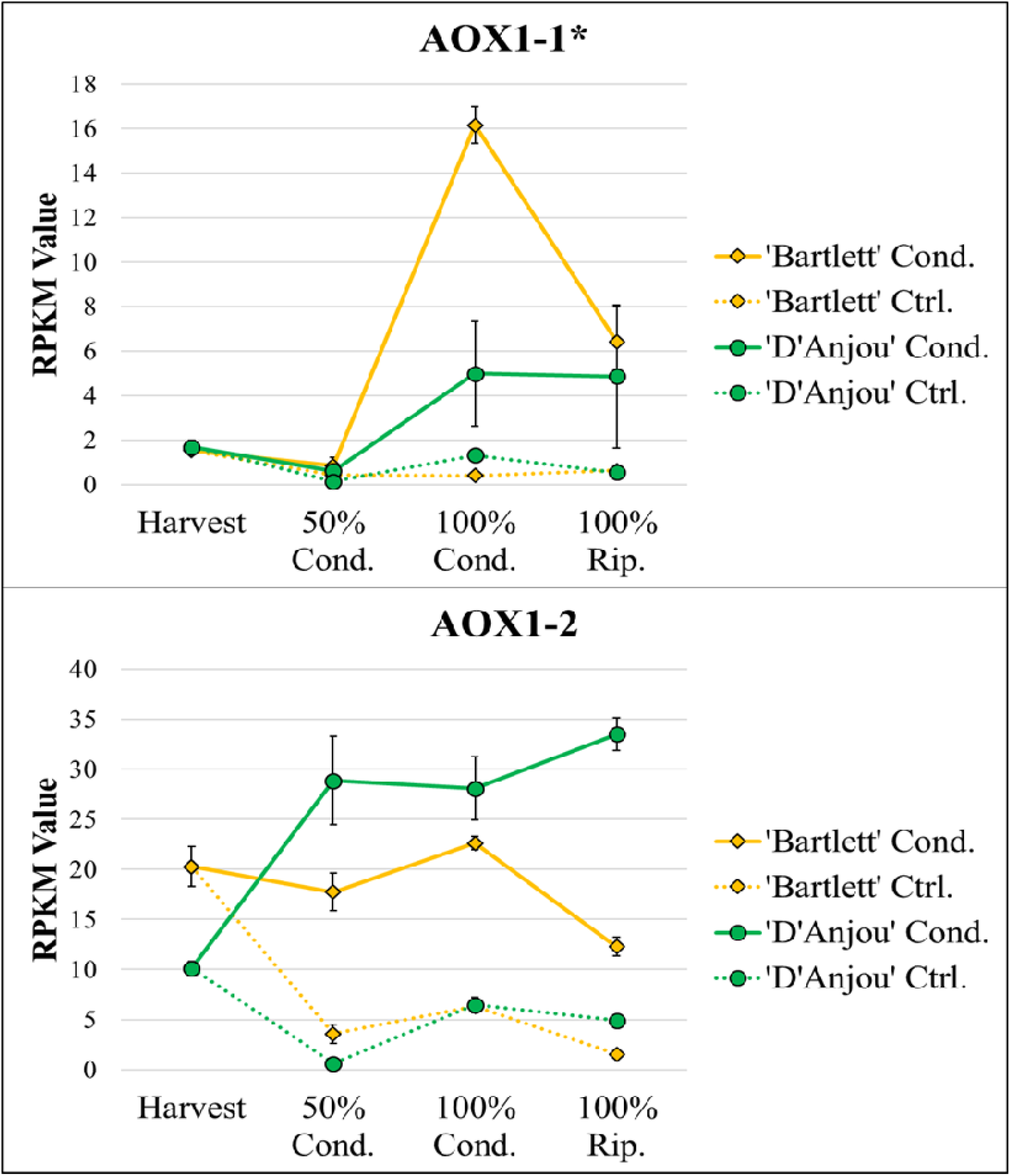
Two homologues of mitochondrial *AOX1* were found to be differentially expressed (p>0.05). Asterisk indicates significant differential expression over time in conditioned ‘Bartlett’, but not in conditioned ‘D’Anjou. Significant linear and quadratic trends (R>0.8) displayed by genes can be seen in Table 1.

**Figure 2.**
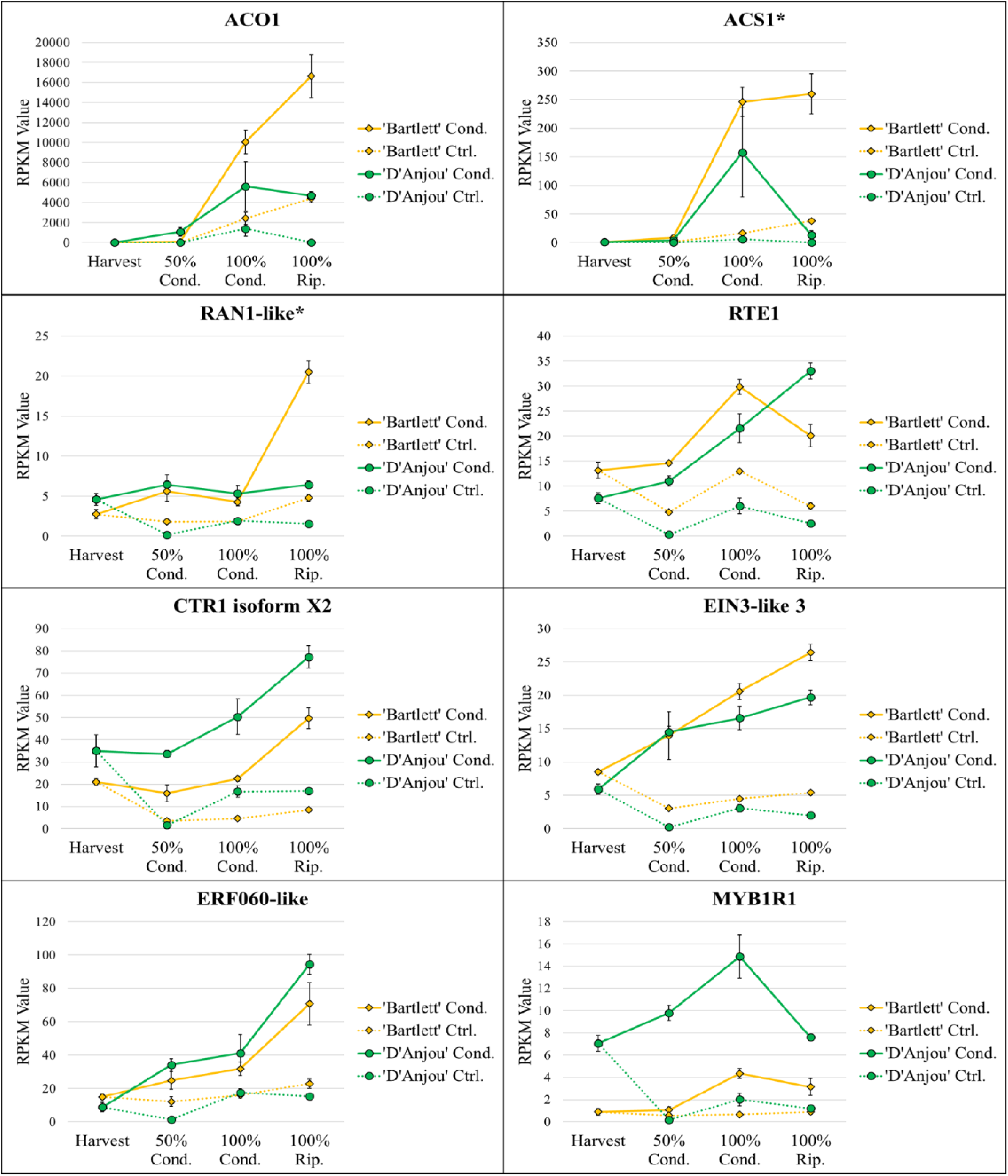
Transcript abundance for differentially expressed ethylene-associated contigs. Asterisk indicates significant differential expression over time in conditioned ‘Bartlett’ but not conditioned ‘D’Anjou’. Significant linear and quadratic trends (R>0.8) displayed by genes can be seen in Table 1.

### Alternative Oxidase

Confirming previous observation, pre-climacteric activation of alternative respiratory pathway transcription was observed during conditioning. Two DECs corresponding to mitochondrial ubiquinol oxidases, homologs of *AOX1*, displayed an increase in expression trend consistent with a preceding report where pre-climacteric increase in *AOX1* was observed as the fruit completed its conditioning requirement [26] (Figure 1). *AOX1* gene expression has been reported in many frwith *AOX* isofouit systems, however mostly at climacteric or post-climacteric stages, rms displaying responses to a broad range of stresses. The different expression patterns of the two *AOX* contigs corresponding to ‘D’Anjou’ and ‘Bartlett’ suggests that *AOX* isoforms differentially regulate responses in different genetic backgrounds. The variable actions of AOX homologues on biological processes has been previously observed in Arabidopsis and tomato [32, 33]. Knock-down of *AOX* in tomato delayed ripening, indicating a regulatory role of AOX in fundamental processes like ethylene response. Furthermore, overexpression of *AOX* in tomato alleviated some of the inhibitory effects of 1-MCP on ripening [34]. In European pear, respiratory partitioning into the alternative pathway may impact S2 ethylene biosynthesis, the climacteric respiration peak, and consequent ripening-related trait development, independent of prior ethylene sensitivity [26, 34, 35].

### Ethylene

For both cultivars, abundance of many transcripts associated with ethylene biosynthesis, perception, and signaling increased throughout the conditioning and ripening period in agreement with similar previous studies [22, 25]. *ACO1* and *ACS1*, were significantly differentially expressed and increased in expression throughout the duration of the conditioning time course in both ‘D’Anjou’ and ‘Bartlett’, as did *RAN1*, which delivers a copper ion that is necessary for ethylene to bind to its receptors [10, 36–38].

Of particular interest in this study was expression pattern of ethylene repressors in response to cold. Consistent with results of recent studies in pear, a *MYB1R* transcription factor, a repressor of ethylene and ripening responses, decreased in expression once ripening competency was reached [21, 39] (Figure 2). Furthermore, brassinosteroid-associated *Brassinazole-Resistant 1* (*BZR1*) and chromatin modification-associated *Multicopy Suppressor of IRA4* (*MSI4*), which have been shown to repress ethylene responses in banana and tomato, respectively, displayed different expression trends in the two cultivars. *BZR1* increased over time in ‘D’Anjou’ and decreased over time in ‘Bartlett’ [40] (Supplementary File 3). These observations may provide further insight into the cultivar specific nature of ripening in pear, as ‘D’Anjou’ is known to be inherently more recalcitrant to ripening [41]. The different ripening trajectories apparent from analysis in this study are congruent with the different vectors followed by the two cultivars in the recent study, which used NMDS analysis to evaluate the relationship between ripening and genes related to the process [26]. The different expression patterns of ethylene signaling genes during ripening suggest a more pronounced ethylene response in the ‘Bartlett’ cultivar, which may be associated with the shorter conditioning time, but perhaps a more complex and a different system of regulation in ‘D’Anjou’.

### Abscisic Acid (ABA)

ABA is well-established as a key regulator of timing of endodormancy release in both model and non-model organisms, such as pear [42, 43]. ABA-related DECs that displayed a similar increase in expression patterns over time for both cultivars included *9-cis-epoxycarotenoid dioxygenase* (*NCED1*), which catalyzes the first step in ABA biosynthesis and regulates some genes associated with cell wall degradation during ripening [44]; *abscisic acid 8’-hydroxylase 2* (*CYP707A*), which is important for regulating seed dormancy and germination in Arabidopsis, accumulating over the course of seed maturation and resulting in the breakdown of ABA [45] (Figure 3). CYP707A has also been shown to inhibit the expression of *NCED-like* genes, thereby reducing ABA biosynthesis in strawberry and tomato [46]. *Abscisic-Aldehyde Oxidase* (*AAO*) related transcript displayed variable expression between the two cultivars (Figure 3). The concomitant increase in transcripts encoding these antagonistic enzymes suggests that increased ABA synthesis is paralleled by a simultaneous increase in the ABA degradation in a tug-of-war between endodormancy maintenance and release, where the latter is favored only when sufficient chilling has occurred.

**Figure 3.**
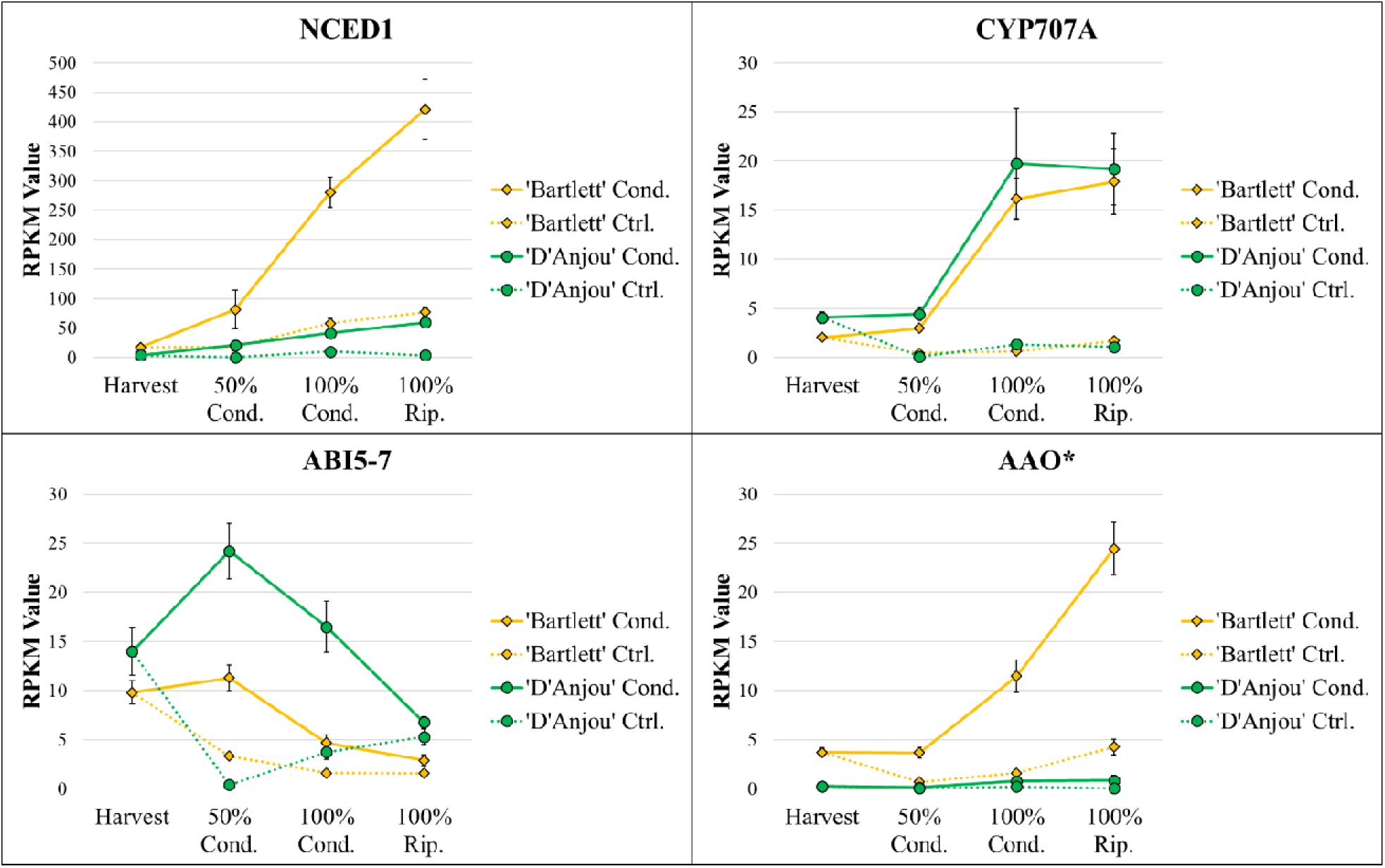
Transcript abundance for differentially expressed ABA-associated contigs (p<0.05). Asterisk indicates significant differential expression over time in conditioned ‘Bartlett’, but not in conditioned ‘D’Anjou’. Significant linear and quadratic trends (R>0.8) displayed by genes can be seen in Table 1.

In contrast to DECs displaying a continual increase in expression over time, *Abscisic Acid Insensitive 5* (*ABI5*) displayed decreased expression in both cultivars during the second half of conditioning and ripening period. This gene is a negative regulator of flowering in Arabidopsis [47] and therefore may play a similar role in negative regulation of other endodormancy-associated processes including fruit ripening.

### Sulfur metabolism

Sulfur containing compounds, including hydrogen sulfide (H_2_S), enhance alternative pathway respiration and inhibit ROS production in fruits [48–51]. Such compounds have also been [50, 51]used in fruit processing as a preservation strategy to reduce oxidative browning [52], and low dose applications of H_2_S elicit a pronounced ripening response in pear fruit [53]. The sulfur metabolism genes, *ATP sulfurylase* and *ATP sulfurylase 2*, were highly expressed only in ‘Bartlett’, with transcript abundance increasing over the course of conditioning and ripening (Supplementary File 4). It is possible, given these results, that sulfur metabolism genes play a role in ripening in a cultivar specific manner.

### Cold and temperature stress-induced processes

In addition to phytohormone-associated genes, several genes and gene families that have previously been implicated in cold-induced endodormancy release and vernalization processes displayed similar expression patterns over time. Those increasing continually during cold conditioning for both cultivars included *Early Flowering 3* (*EF3*), which has been shown to maintains the circadian clock in a temperature-dependent manner in barley and Arabidopsis [54] (Figure 4). Upregulation of this regulatory gene in pear fruit exposed to chilling may result in induction of genes associated with release of endodormancy. *Polycomb group embryonic flower 2-like isoform x1* (*EMF2*) decreased continually in expression during conditioning and ripening in ‘Bartlett’ (Supplementary File 5). In Arabidopsis, loss of function of EMF2 causes direct initiation of flowering, causing a bypass of vegetative shoot growth [55]. The isoform present in ‘Bartlett’ may play a similar role in modulating initiation of ripening.

**Figure 4.**
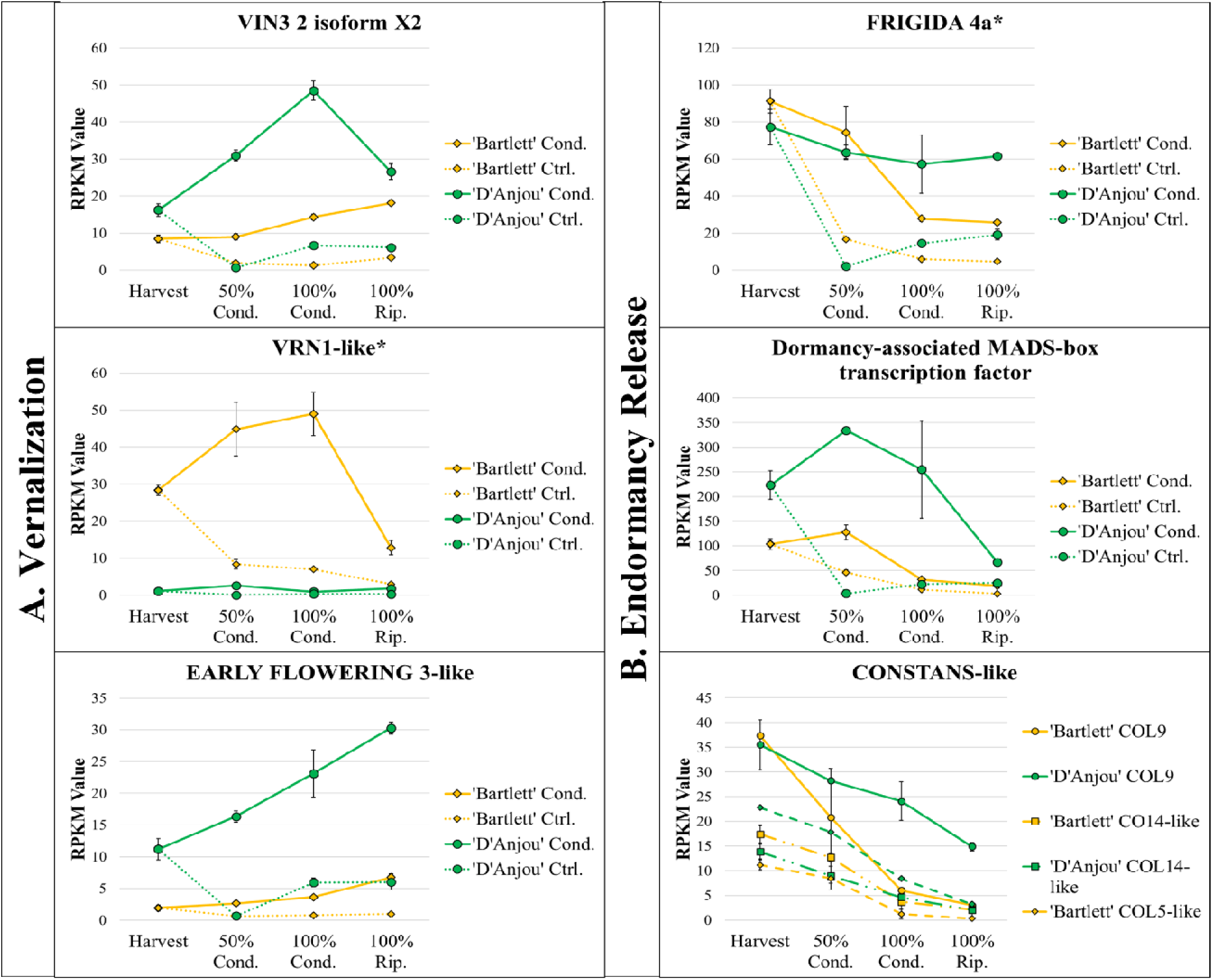
Differentially expressed vernalization-associated genes *VIN3* and *VRN1* and cold responsive *ERF3-like* (p>0.05) (column A) and differentially expressed endodormancy release-repressing genes FRIGIDA 4a, dormancy-associated MADS-box transcription factor, and CONSTANS-like over conditioning time course (p>0.05) (column B). Asterisks indicate significant differential expression over time in conditioned ‘Bartlett’ but not in conditioned ‘D’Anjou’. Significant linear and quadratic trends (R>0.8) displayed by genes can be seen in Table 1.

Furthermore, a DEC corresponding to a BRCA1 homolog increased during conditioning. The *BRCA1* gene has homologues in humans, and BRCA mutations are most often associated with increased cancer susceptibility; however, the primary function of the gene is DNA damage repair and chromatin remodeling, and BRCA1 has been shown to play a similar reparatory role in plants [56, 57]. It is possible that BRCA1 in fruit, might play a role in mediating temperature-induced stress damage to DNA during cold conditioning. In addition to *BRCA1*, *Next-to-BRCA* (*NBR1*) displayed increasing expression during conditioning and ripening for both cultivars (Supplementary File 6). NBR1 plays a role in heat stress tolerance in Arabidopsis [58], although little work has been done to study its effects in other plants.

The vernalization-associated gene *VRN1*, has been characterized with regards to flowering time in both model and non-model species, specifically in the context of cold [15, 59]. In cereal grains transcriptional activation of *VRN1* after prolonged chilling results in accelerated flowering [60]. Interestingly, *VRN1* was significantly differentially expressed over time in ‘Bartlett’ during the accumulation of chilling hours, while expression levels remained low in ‘D’Anjou’ throughout the time course (Figure 4). In ‘Bartlett’ the observed patterns of expression of *VRN1* during conditioning are consistent with a previously described model in wheat, in which increased accumulation of *VRN1* transcripts correlates with a decreased repression of endodormancy release, primarily via repression of FLC-like genes and other developmental repressors [61, 62].

Another vernalization-associated gene, *VIN3* isoform x1, which is associated with temperature-mediated epigenetic regulation of endodormancy repressors [63], displayed increasing expression in both cultivars during the cold conditioning period (Figure 4). As VIN3 and VRN1 are both cold-induced repressors of endodormancy release and are expressed differently in ‘D’Anjou’ and ‘Bartlett’, the opposite, genotypic-specific expression of these two DECs suggests that cold induced ripening might occur via two different vernalization-associated pathways, and may influence the duration of cold-requirement in different pear cultivars.

Working in an antagonistic manner to VRN1 and VIN3, which downregulate repressors of endodormancy release, are *FRIGIDA 4* and *CONSTANS-like* genes. Increased expression and activity of *FRIGIDA 4* results in increased activity of repressors of endodormancy release, such as *Flowering Locus C* (*FLC*), in Arabidopsis [64] and blueberry [65]. *FRIGIDA 4* decreased significantly in expression throughout conditioning in ‘Bartlett’, suggesting that its downregulation may correspond to decreased activation of ripening repressors in a role homologous to regulation of flowering time (Figure 4).

In addition to *FRIGIDA 4*, overexpression of *CONSTANS-like 9* has been shown to delay flowering and regulate the circadian clock by repressing *CONSTANS* and Flowering Locus T (*FT)* gene expression in photoperiod sensitive plants, like Arabidopsis [66]. Recently, however, *CONSTANS-like* gene families have been shown to display distinct, tissue-specific patterns of expression in banana fruit and pulp, in addition to other tissues [67], suggesting that *CONSTANS-like* gene family members play a role not only in flower development, but also, fruit development and senescence. *CONSTANS-like* 9, 5, 6, 13, and 14 were all differentially expressed. Overall, the *CONSTANS-like* genes displayed a similar decreasing expression trend in both cultivars throughout the conditioning period (Figure 4). As with *FRIGIDA 4*, this decrease suggests that the repressive role of these genes with regards to process of endodormancy release is downregulated during cold conditioning, thereby promoting ripening.

### Functional Enrichment Analysis

Shared overrepresented terms included ‘cold acclimation’, dormancy-related (‘embryo development ending in seed dormancy’, ‘flowering’), hormone signaling (‘ethylene-activated signaling pathway’, ‘ABA-activated signaling pathway’, ‘auxin-activated signaling pathway’), hormone biosynthesis (‘jasmonic acid biosynthetic process’, ‘salicylic acid biosynthetic process’, ‘brassinosteroid biosynthetic process’), and respiration-associated processes (‘ATP synthesis coupled proton transport’, ‘electron transfer activity’) (Figure 5). The shared overrepresentation of cold- and dormancy-related GO terms lends support to mediation of cold-induced ripening responses by vernalization-associated genes, in conjunction with downregulation of repressors of endodormancy release. Additionally, the presence of many enriched phytohormone-related ontologies lends support to the concept of interacting networks of phytohormonal crosstalk that serve to mediate ripening and mitigate chilling injury [22, 29, 68]. This lends support to the importance of auxin biosynthesis and metabolism during the ripening process in *Pyrus*. Finally, enriched ontologies associated with mitochondrial respiration, which implicate a high rate of ATP production, provide further evidence that AOX1 alternative respiratory activity is needed to alleviate some of the stress on the cytochrome respiratory pathway during ripening [69]. While shared enriched GO terms lend insight into conserved biological basis for cold-conditioning mediated ripening, enriched ontologies unique to ‘D’Anjou’ or ‘Bartlett’ provide information regarding the cultivar-specific ripening responses.

**Figure 5.**
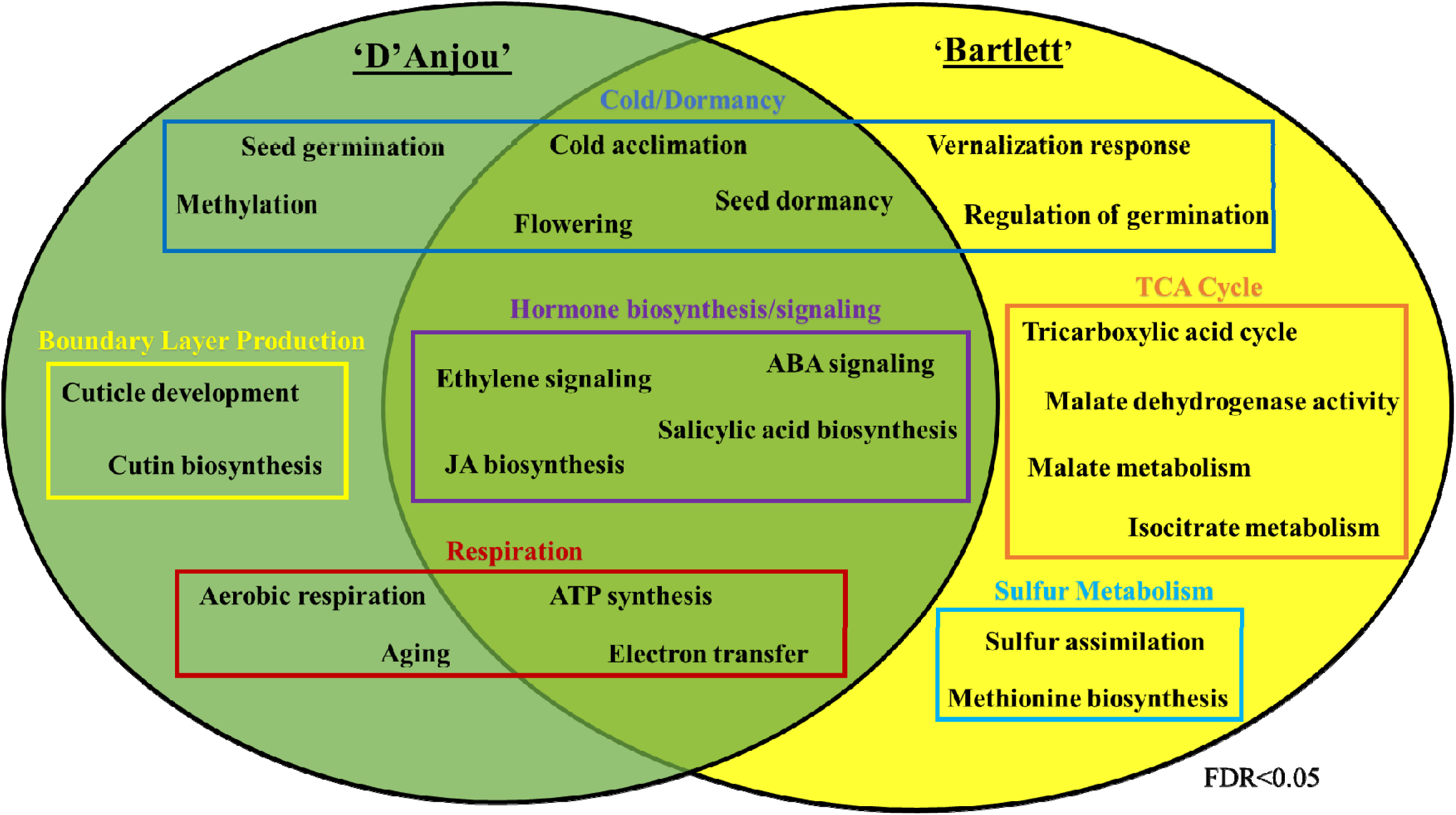
Selection of shared and unique overrepresented GO terms shared between ‘D’Anjou’ and ‘Bartlett’ cultivars that were identified using the OmicsBox enrichment analysis feature. Additional values can be seen in Supplementary File 7.

‘D’Anjou’ pears are genetically programmed to require a longer conditioning time to ripen than ‘Bartlett’ pears (60 versus 15 days of conditioning at 0°C). Overrepresented GO terms unique to ‘D’Anjou’ associated with chilling-induced endodormancy release included ‘seed germination’ and ‘methylation’, while for ‘Bartlett’ such terms included ‘vernalization response’, ‘regulation of seed germination’ (Figure 5). Regulation of vernalization sensitive genes, including homologs of those governing timing of seedling germination in many crops, is highly dependent on methylation status and other epigenetic modifications, suggesting that such genes may play a larger role in the ripening of ‘D’Anjou’. Further investigation is needed of the effects of external abiotic factors like chilling on the epigenome of fruit undergoing developmental transitions such as a shift to ripening. The overrepresentation of terms associated with respiration and senescence are expected, as such processes are characteristic of ripening and the terminal stages of fruit development. In ‘D’Anjou’, the terms ‘aerobic respiration’ and ‘aging’ were overrepresented, while in ‘Bartlett’, TCA cycle-associated terms (‘tricarboxylic acid cycle’, ‘malate metabolic process’, ‘malate dehydrogenase activity’, ‘isocitrate metabolic process’) were overrepresented (Figure 5). Differential overrepresentation of aerobic respiration and TCA cycle metabolism GO terms in the two cultivars suggest that these processes are under cultivar-specific regulation. Interestingly, enrichment of terms associated with production of protective boundary layers (‘cutin biosynthetic process’, ‘cuticle development’) may represent a genetically programmed stress management strategy considering long conditioning requirements. Development of such barriers could mitigate the occurrence of chilling injury while ‘D’Anjou’ fruits accumulate the required chilling hours (Figure 5. In ‘Bartlett’, ‘sulfur assimilation’ and ‘methionine metabolic process’ were enriched. This is interesting because the Yang cycle, which recycles the sulfur containing amino acid methionine, also feeds into the production of the ethylene biosynthetic precursor ACC. Increased sulfur metabolic capacity in ‘Bartlett’ may in turn correspond to higher methionine cycling capacity, and therefore production of ethylene for this cultivar. Based on this finding, and that of the DE analysis, in which ATP sulfurylases were highly expressed during conditioning and ripening in ‘Bartlett,’ it is possible that cold conditioning directly or indirectly induces sulfur metabolism, thereby inducing ethylene biosynthesis and downstream processes. This is the case for soybean, in which ATP-sulfurylase is induced by cold and catalyzes activation of sulfate [70, 71]. The GO analysis results implicate cold temperature induction of numerous metabolic pathways, many of which operate upstream or independently of ethylene. This observation aligns with a recent study in cold conditioned pear fruit that demonstrated that LT induces expression of both ethylene-dependent and independent genes affecting ripening [25]. Complete ontology results can be found in Supplementary File 7.

To summarize, the ontology enrichment results provide a global overview with regards to some of the overarching processes responsible for cold-induced, ripening induction in pear. These results lend credibility to the role of vernalization-associated genes, and their potential role in influencing the duration of cold required for conditioning for fruit ripening.

### Conclusion

In this study, time-course differential expression analysis, functional annotation and GO enrichment methods were used to identify candidate genes and gene networks associated with the chilling requirement for ripening in pear. The results agree with previously reported expression patterns of known ripening-related genes during achievement of ripening competency, specifically genes associated with ethylene biosynthesis and phytohormonal crosstalk. A novel outcome of this study was that differentially expressed cold-responsive, vernalization-associated genes may play a role in the ripening of European pear. While described in other systems, these genes have not yet been characterized with respect to their role in climacteric ripening.

Notably, *AOX* expression results are consistent with our recent work in pear, providing support for the possible role of cold-induced AOX activity in the achievement of ripening competency. AOX has been described previously in the context of cold stress response and ROS mediation, and more recently in pre-climacteric S2-S2 transitionary phase [26, 53, 69]. Its expression and activity may be linked to or activated in conjunction with vernalization-associated genes via ROS as a response to cold temperatures (Figure 6). Further studies are needed to elucidate the precise connections, but it is clear based on expression data that these transcripts share a similar response to cold conditioning in pear fruit.

**Figure 6.**
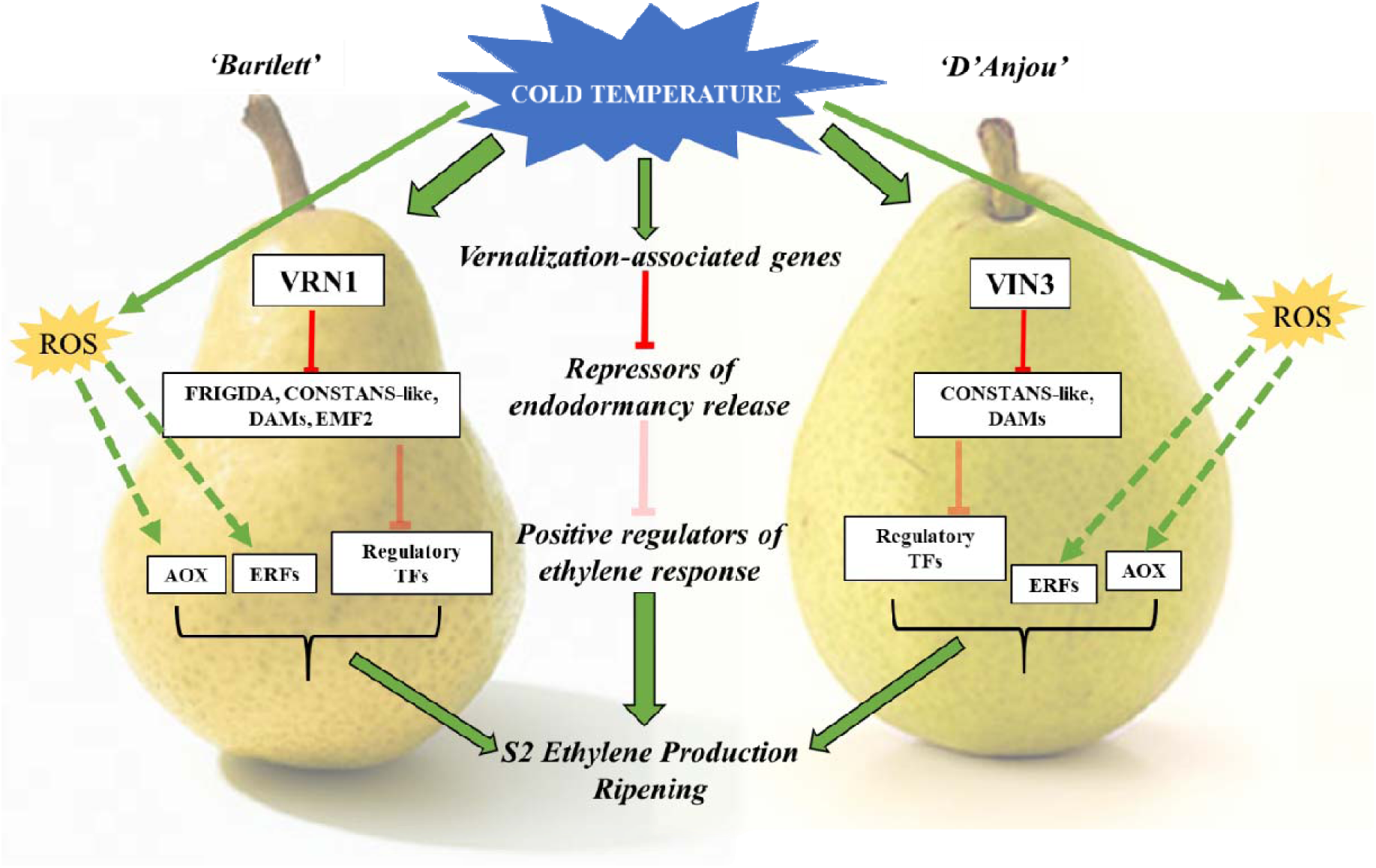
Model describing a possible mechanism by which vernalization-associated genes may mediate cold-induced ripening. Cold temperature stimulates VRN1/VIN3. Vernalization genes, in turn inhibit repressors of endodormancy release—FRIGIDA, CONSTANS-like, DAMs, EMF2. Downregulation of these repressors allows for transcriptional activation of ethylene response. Cold also triggers ROS signaling, leading to activation of ERFs [72] and AOX [27]. S2 ethylene production is triggered and ripening commences.

Based on these findings, the mechanism by which vernalization-associated genes may mediate cold-induced ripening may manifest as follows: Cold temperature stimulates VIN3 or VRN1 in a cultivar dependent manner. These, in turn inhibit fruit-tissue specific repressors of endodormancy release (CONSTANS-like 9, FRIGIDA 4, dormancy-associated MADS-box genes, and ERF2). Inhibition of repressors of endodormancy release, such as BZR1 and MSI4 and ABA precursors, via VRN1 and VIN3 pathways allows for activation of ripening-specific transcription factors, such as MYB1R1 and others, which may regulate autocatalytic ethylene production during conditioning [22]. ROS induced by cold temperatures may concurrently serve to activate AOX and ethylene response factors [27, 72]. The normal ripening climacteric, characterized by the conversion of ACC to ethylene by ACS, commences following transcription factor-mediated activation. Ethylene biosynthesis and response results in activation of downstream ripening processes (Figure 6).

Results of this study provide new information with regards to vernalization and cold response-associated genes that are differentially expressed over time during conditioning and subsequent ripening. Many of the genes which have been identified as potential regulators of chilling-induced ripening in pear fruit represent members of diverse gene families. Furthermore, several studies have previously indicated the diversification and neofunctionalization of *VRN*, *VIN*, *CONSTANS-like* gene families among others, in a multitude of plant tissues, including roots, shoots, leaves, apical meristems, buds, and flowers [67, 73]. Here we provide evidence suggesting that members of these gene families have diversified to play similar roles in chilling-dependent fruit ripening. These findings lend support to the idea of chilling-induced ripening as a process of endodormancy release that might explain the underlying basis of different chilling requirement across different cultivars.

## Materials and Methods

### Experimental Design

The experimental design was as previously reported [26]. Briefly, ‘Bartlett’ and ‘D’Anjou’ pear fruit were obtained from Blue Star Growers (Cashmere, Washington). During the time between harvest and acquisition (5 days), the pear fruit was maintained in temporary storage at 1°C. ‘Bartlett’ fruit had a mean firmness of 76.2 N, and 13.40 °Brix and ‘D’Anjou’ fruit had a mean firmness of 53.5 N, and 12.66 °Brix at initiation of the experiment. Ripening of ‘Bartlett’ requires 15 days of cold conditioning, while ‘D’Anjou’ typically requires 60 days of −1°C to attain ripening competency [9, 10]. The duration of cold conditioning, however, is reduced when conditioning temperatures are increased to 10^°^C [8]. Pears were divided into equal replicate groups and then placed into storage at 10°C for conditioning [8]. After the conditioning period (Figure 7), the fruit was transferred to 180-liter flow-through respiration chambers held at 20°C for seven days. The flow rate of the chambers was maintained at 5.0 ml/min with compressed air. Fruit was evaluated at four physiological time points: at 0% conditioned, 50% conditioned, and 100% conditioned and 100% ripened, which comprised 7 days after completion of conditioning. The pears that were allowed to accumulate the required chilling hours for ripening were transferred to 20°C and were then sampled at the 100% ripened time point (Figure 7). These time points were similar to previously utilized physiological stages determined during conditioning [26].

**Figure 7.**
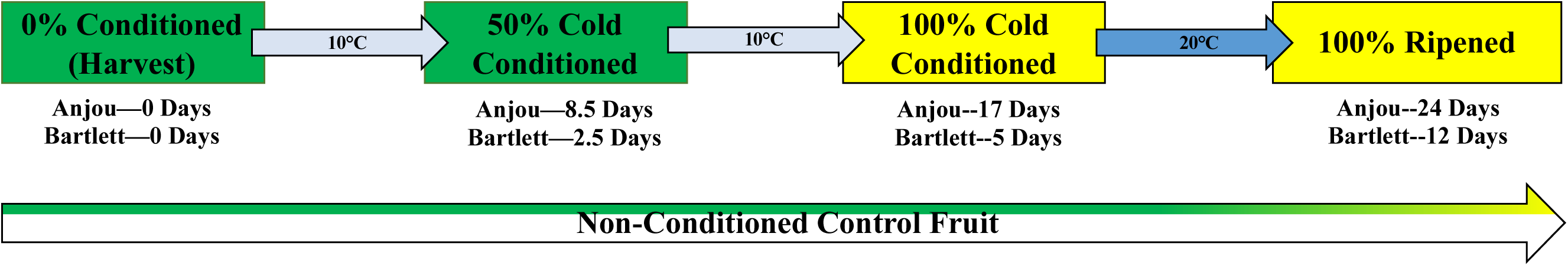
Conditioning time course schematic used for ‘D’Anjou’ and ‘Bartlett’. Conditioning times were determined based on the established, cultivar-specific chilling conditions required for achievement of competency and ripening.

### Fruit firmness measurements and tissue sampling

Firmness was measured for 10 replicate fruit at each sampling time point. A GS-14 Fruit Texture Analyzer (GÜSS Instruments, South Africa) equipped with an 8.0 mm probe set at 5.0 mm flesh penetration was used to measure firmness at two equidistant points around the equatorial region of each fruit after removal of the peel. Firmness data were assessed using ANOVA, following the statistical approaches described previously [74, 75].

### RNA Extraction

Peel tissue used in this study was the same as that used for the qRT-PCR study, published recently [26]. Briefly, peel tissue was obtained from a 1 cm wide equatorial region of 3 randomly sampled fruit of each cultivar and at each conditioning time point, flash frozen in liquid nitrogen, pooled for each treatment/time point, and then ground using a SPEX Freezer/Mill 6870 (Metuchen, NJ USA). Total RNA was extracted from pulverized ‘D’Anjou’ and ‘Bartlett’ peel tissue for each of the four technical replicates at 0% conditioned, 50% conditioned, 100% conditioned, and 100% ripened time points following the methods of [76]. Contaminating genomic DNA was removed with DNaseI per manufacturer instructions (NEB, Ipswich, MA USA). RNA was quality checked using a denaturing gel and BioAnalyzer 2100 (Agilent, CA, USA) and was quantified using Nanodrop 2000 spectrophotometer (Thermo Scientific, Waltham, MA, USA).

### Illumina Sequencing

cDNA libraries were qualified and quantified using a Life Technologies Qubit Fluorometer (Carlsbad, CA) as well as an Agilent 2100 Bioanalyzer (Santa Clara, CA). The cDNA libraries prepared from the extracted RNA were sequenced on an Illumina Hi Seq 4000 platform as 2×100 paired end reads. The Illumina TruSeq RNA Sample Preparation v2 kit (San Diego, CA) was used to generate the final library molecules, and the Ailine Biosciences’ (Woburn, MA) DNA SizeSelector-I bead protocol was used to filter for library molecules of >450 base pairs.

### Transcriptome Assembly

Transcriptome assembly was performed as reported previously [77, 78]. The 2×100 paired end fastq files generated using Illumina HiSeq 2000 were input into the CLC Bio Genomics Workbench (ver 6.0.1) (Aarhus, Denmark) for pre-processing and assembly. The CLC Create Sequencing QC report tool was used to assess quality. The CLC Trim Sequence process was used to trim quality scores with a limit of 0.001, corresponding to a Phred value of 30. Ambiguous nucleotides were trimmed, and the 13 5’ terminal nucleotides removed. Reads below length 34 were discarded. Overlapping pairs were merged using the ‘Merge Overlapping Pairs’ tool, and a subsequent de novo assembly was performed with all datasets. Parameters used in the assembly are as follows: Map reads back to contigs = TRUE, Mismatch cost = 2, Insertion cost = 3, Deletion cost = 0.4, Similarity Fraction = 0.95, Global Alignment = TRUE, Minimum contig length = 200, Update contigs = true, Auto-detect paired distances = TRUE, Create list of unmapped reads = TRUE, Perform scaffolding = TRUE. The de novo assembly resulted in the production of 140,077 contiguous sequences (contigs). Contigs with less than 2× coverage and those less than 200bp in length were eliminated. For each individual dataset (treatment/replicate) the original, non-trimmed reads were mapped back to the master assembly subset. Default parameters were used, except for the length fraction, which was set to 0.5, and the similarity fraction, which was set to 0.9. Mapping resulted in the generation of individual treatment sample reads per contig. The master transcriptome was exported as a fasta file for functional annotation and the read counts for each dataset were exported for normalization with the Reads Per Kilobase per Million reads (RPKM) method [79].

### Functional Annotation with Blast2GO

The master transcriptome fasta produced from the Illumina assembly was imported into OmicsBox (BioBam Bioinformatics S.L., Valencia, Spain) for functional annotation of expressed contigs using the Blast2GO feature [80]. Contig sequences were identified by a blastx alignment against the NCBI ‘Viridiplantae’ database with and e-value specification of 10.0E-3. Gene ontology (GO) annotation was assigned using the ‘Mapping’ and ‘Annotation’ features. Expression analysis was limited to the consensus sequence for each contig, and therefore in this paper we do not distinguish between specific alleles, highly similar gene family members. This is due to assembler constraints [81].

### Differential Expression Analysis

An Excel file was prepared containing ‘D’Anjou’ and ‘Bartlett’ RPKM data for each contig, treatment, and replicate. The data was imported into OmicsBox as a count table for differential expression analysis, which employs the maSigPro R package [82]. An additional experimental design matrix was imported which dictated the number of time points and replicates (Supplementary File 8). The level of FDR control was set to 0.05, resulting in identification of significant genes. A stepwise regression was employed to model the data and then generate a list of all genes displaying significant linear or quadratic trends over the cold conditioning time course (R>0.8) [83] (Supplementary File 2).

### GO Enrichment Analysis

OmicsBox gene ontology (GO) enrichment analysis utilizing the Fisher’s Exact Test was employed [80]. Due to many enriched GO terms, the resulting terms were reduced to only the most specific ontologies (p<0.00001). Ontologies shared between ‘D’ Anjou’ and ‘Bartlett’ and unique to each cultivar were identified (Supplementary File 7).

### qRT-PCR validation

qRT-PCR was performed as reported earlier [26]. Briefly, RNA samples were treated with DNAseI to eliminate any DNA contamination according to the manufacturer’s methods (NEB, Ipswich, MA USA), prior to cDNA synthesis. RNA concentration was determined for each sample using a Nanodrop ND-8000 (ThermoFisher, MA, USA). RNA quality was verified using a denaturing gel and BioAnalyzer 2100 (Agilent, CA USA). For each sample, 500 ng of total RNA was used to generate first strand cDNA using the Invitrogen VILO kit (Life Technologies, Carlsbad, CA USA). Each cDNA preparation was quantified, and the mean concentration calculated from eight replicate quantification measurements, recorded using a NanoDrop8000 (Thermo Fisher Scientific, Waltham, MA). The samples were diluted to a final concentration of 50 ng/uL. Initial qRT-PCR technical replicate reactions were prepared for each of the 90 selected genes using the iTaq Universal SYBR Green Supermix (BioRad, Hercules, CA). Primers for quantitative reverse transcriptase PCR (qRT-PCR) were designed from Pyrus ESTs or sequences derived from *Malus* × *domestica* transcripts among various hormonal and environmental signaling pathways. 500ng RNA for each sample (same as used for RNAseq) was used to generate 1st strand cDNA using the Invitrogen VILO kit (Life Technologies, Carlsbad, CA USA). cDNA preparations were then diluted to 50ng/uL. qRT-PCR technical replicate reactions were prepared for each of the genes using the iTAq Universal SYBR Green Supermix with ROX reference dye (BioRad, Hercules, CA) per the manufacturer’s protocols with 100ng of template cDNA. In a Strategene MX3005P, the following thermocycle profile was used: 95°C initial disassociation for 150s followed by 50 amplification cycles (95°C for 30s, 60°C for 30s, and 72°C for 30s) and a final, single cycle phase to generate a dissociation curve (95°C for 150s, 95°C for 30s, and 60°C for 30s).Using the LinRegPCR tool, we calculated the Cq values for each reaction [84, 85] (Supplementary File 9).

## Supporting information

S1_PearTranscriptome_Annotated text

S2_Anj_Bart_Cold_RPKMs

S3_EthyleneReg

S4_Sulfur_DECs

S5_EMB2

S6_BRCA_DECs

S7_EnrichedGOs_All

S8_maSigPro_Data_and_Designs_All

S9_qRT-PCR_validation_all

## Acknowledgements

The authors thank Blue Bird Growers (Peshastin, WA) and Blue Star Growers (Cashmere, WA) for providing pears for conditioning experiments and to Scott Mattinson for assistance in maintenance of the experimental infrastructure. Work in the Dhingra lab was supported in part by Washington State University Agriculture Center Research Hatch Grant WNP00011 and grant funding from Pear Bureau NW to AD. SLH acknowledges the support received from ARCS Seattle Chapter and National Institutes of Health/National Institute of General Medical Sciences through an institutional training grant award T32-GM008336. The contents of this work are solely the responsibility of the authors and do not necessarily represent the official views of the NIGMS or NIH.

## SUPPLEMENTARY FILES

**Supplementary File 1.** Annotated master assembly fasta for cold conditioned ‘D’Anjou’ and ‘Bartlett’ pear fruit.

**Supplementary File 2.** Mean RPKM values, standard error, and time course differential expression information for cold conditioned ‘D’Anjou’ and ‘Bartlett’ pear fruit.

**Supplementary File 3.** Differential expression graphs for of ethylene regulatory genes *BZR1* and *MSI4*

**Supplementary File 4.** Differential expression graphs for sulfur metabolism-associated genes.

**Supplementary File 5.** Differential expression graph for *Polycomb group embryonic flower 2-like isoform x1* (*EMF2*).

**Supplementary File 6.** Differential expression graphs for BRCA1 and Next-to-BRCA1 genes.

**Supplementary File 7.** All shared and unique enriched gene ontologies for cold conditioned ‘D’Anjou’ and ‘Bartlett’ pear fruit.

**Supplementary File 8.** RPKM data files and experimental design files used for MaSigPro time-series differential expression analysis in OmicsBox.

**Supplementary File 9.** Quantitative RT-PCR validation with calculated expression values.

